# A brain-specific microRNA, miR-1000, regulates lipid homeostasis via Neuropeptide-like precursor 1 in *Drosophila melanogaster*

**DOI:** 10.1101/2025.07.22.666178

**Authors:** Pushpa Verma, Pruthvi Gowda, Nika N Danial, David Van Vactor

## Abstract

Metabolism requires precise gene regulation to balance energy intake and expenditure for an organism’s well-being, with misregulation often leading to metabolic syndromes. This study reveals that the brain-specific microRNA *miR-1000* regulates fat storage by controlling the expression of a neuropeptide gene, *Nplp1*. Loss of *miR-1000* increases *Nplp1* expression, leading to higher body weight, increased fat storage, improved survival under food deprivation conditions, and a reduced overall lifespan in *Drosophila*. We further show that *miR-1000* promotes fat storage upon feeding by regulating TAG synthesis and storage in lipid droplets, thereby playing a crucial role in metabolic regulation.

## Introduction

Metabolic homeostasis is a highly complex process that demands a delicate equilibrium between energy storage and expenditure across multiple organ systems. Misregulation of metabolism often leads to disease conditions such as obesity, diabetes, cardiovascular diseases, or elevated triglycerides and cholesterol, which contribute significantly to global mortality. While considerable focus has been placed on glucose and carbohydrate metabolism, disruptions in lipid metabolism are equally implicated in the pathogenesis of metabolic syndromes and diseases, including obesity and diabetes (McGarry 1992). Lipids provide more energy per gram than carbohydrates and can be readily mobilized to support organisms during food deprivation. Notably, lipid storage correlates positively with increased survival during starvation (Ballard et al. 2008). Maintaining a precise balance between lipid storage and mobilization is vital for metabolic health. Therefore, understanding the regulatory mechanisms governing lipid metabolism is of paramount importance.

The nervous system plays a pivotal role in metabolic regulation, encompassing processes such as sensory information processing, initiation and cessation of food intake, and control of the gut and other internal organs (Droujinine and Perrimon 2016; Scopelliti et al. 2019). Peripheral organs, including the gut, actively monitor nutrients and provide feedback to the nervous system (Mayer 2011; Steinert and Beglinger 2011; Dickson 2018). Notably, neuropeptides have been identified as key mediators of neuronal control over peripheral organs, functioning either as local synaptic neuromodulators or as endocrine factors with broader systemic effects (Clynen et al. 2010; Nassel and Zandawala 2019). These neuropeptides are typically expressed as pro-peptide precursors, which undergo proteolytic cleavage to produce active peptides capable of mediating inter-organ communication. *Drosophila* serves as an invaluable model for investigating neuropeptide biology due to its advanced molecular genetics tools and imaging techniques, enabling detailed analysis of cell-specific neuropeptide functions and receptor interactions. Approximately 50 genes in *Drosophila* encode neuropeptide precursors, with the expression patterns of 30 neuropeptide genes, including *DILPs* (*Drosophila Insulin-like peptides*), mapped within the central nervous system (CNS) (reviewed in (Nassel and Zandawala 2019). While the functions of insulin-related neuropeptides along the gut-brain axis are well-documented, the biological roles and regulatory dynamics of many other neuropeptides and their associated neural circuits in maintaining metabolic homeostasis remain to be fully elucidated.

Small 21-23 nucleotide long, non-coding microRNAs (miRNAs) are well-suited as regulators of physiological and adaptive responses that demand rapid and/or localized control over gene expression (He and Hannon 2004; Ebert and Sharp 2012). miRNAs modulate gene expression by inhibiting protein translation and destabilizing mRNA targets containing a “seed sequence” or miRNA Response Element (MRE) capable of base-pairing with the miRNA sequence (Brennecke et al. 2005; Bartel 2009). However, the specificity of this mechanism is determined not only by MRE sequence matching but also by the expression domain overlap of miRNA and target mRNA(s); hence, the same miRNA can act in different ways simultaneously in distinct cell types (Wang and Wang 2006). This property is extensively utilized in the central nervous system (CNS), where coordinated molecular events across multiple cells within a circuit are critical (Cochella and Hobert 2012; McNeill and Van Vactor 2012; Cao et al. 2016). miRNA-mediated regulation provides a versatile and rapid mechanism for controlling target gene expression both locally and globally.

In this study, we identify a novel neural circuit in *Drosophila* that regulates metabolic homeostasis through the conserved CNS-specific microRNA (miRNA), *miR-1000*. We demonstrate that *miR-1000* plays a critical role in controlling body weight, fat storage, survival under nutrient deprivation, and longevity, by targeting the Neuropeptide Like Peptide 1 *(Nplp1)* gene in the central nervous system (CNS). The loss of *miR-1000* leads to increased fat storage and extended survival during nutrient deprivation, driven by elevated *Nplp1* expression. While the biological function of *Nplp1* neuropeptides has remained largely unknown, our findings reveal a significant and novel role for *Nplp1* in maintaining metabolic homeostasis in *Drosophila*. Additionally, we observe dynamic changes in *miR-1000* and *Nplp1* levels during refeeding following starvation, suggesting an adaptive function in metabolic regulation.

## Results and Discussion

### *miR-1000* regulates body weight, fat storage, and survival under nutrient deprivation

*miR-1000* is a conserved *Drosophila* miRNA (paralog of human and mouse *miR-137*) with widespread expression in the CNS. Analysis of the trans heterozygous (*KO1/KO2*) combination of two different *miR-1000* null mutants (*KO1* and *KO2* as described in (Verma et al. 2015) revealed that *miR-1000* mutants are bigger (Fig. 1A) and exhibit increased body weight compared to wild-type (WT) control flies in both males and females (Fig. 1A, S1_A). Biochemical analysis of total glucose and triglycerides (normalized to total protein) showed that *miR-1000* mutants do not affect circulating glucose levels (Fig. 1C) but store significantly higher levels of fat compared to the WT control (Fig. 1D), suggesting that *miR-1000* may regulate fat metabolic efficiency. Consistent with the increased fat reserves, *miR-1000* mutants survived much longer compared to WT control flies upon starvation (Fig. 1E, S1_B), presumably by utilizing these larger fat reserves. Reintroduction of *miR-1000* at its endogenous genomic locus in a *miR-1000* mutant (*KO2*) in *“miR-1000 Rescue”* flies significantly reduced body weight (Fig. 1B, S1_A), fat levels (Fig. 1D), and starvation resistance (Fig. 1E, S1_B) of the *miR-1000* mutants. These findings confirm that *miR-1000* is crucial for maintaining proper metabolic homeostasis of an organism by controlling body mass, fat storage, and the ability to survive food deprivation.

**Figure 1:**
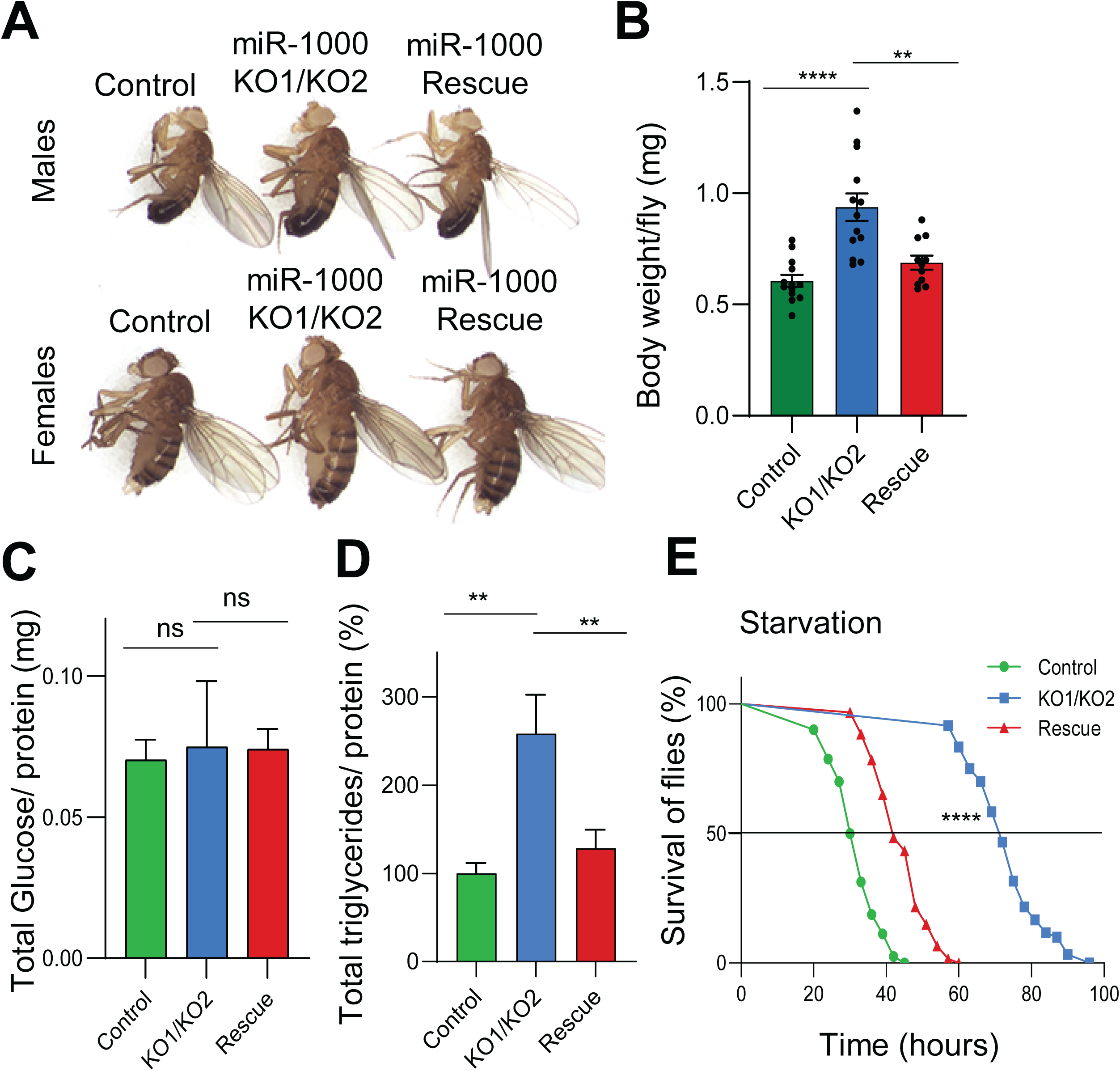
Loss of *miR-1000* causes increased body weight, fat storage, and starvation resistance. A) Adult male (upper panel) and female (lower panel) flies show a bigger size of *miR-1000* mutants (*KO1/KO2*) compared to WT (Wild Type) control and *miR-1000 Rescue* flies. B) Body weight of male flies showing *KO1/KO2* flies are heavier than *miR-1000 Rescue* and WT control. Bars represent an average of at least 11 biological replicates, n=10 flies each, error bars ±SEM. p<0.0001 *KO1/KO2* (n=13) vs WT control (n=13) and p=0.0011 *KO1/KO2* vs *Rescue* (n=11) using one-way ANOVA with Tukey’s multiple comparisons. C) No change in total Glucose levels (normalized to total protein) in *KO1/KO2* flies compared to *miR-1000 Rescue* (p=0.9968) and WT control (p=0.9026) using one-way ANOVA with Tukey’s multiple comparisons. Bars represent an average of at least four biological replicates, n=10 flies each, error bars ±SD D) Increased triglyceride levels (normalized to total protein) in *KO1/KO2* compared to *miR-1000 Rescue* and WT control. Bars represent an average of three biological replicates, n=15 flies each, error bars ±SD. p-value=0.0014 *KO1/KO2* vs WT control and p-value=0.0039 *KO1/KO2* vs *miR-1000 Rescue* using one-way ANOVA with Tukey’s multiple comparisons. E) Percent survival of male flies upon starvation. *KO1/KO2* (n=60) survived longer compared to *miR-1000 Rescue* (n=60) and WT control (n=80). **** p<0.0001 comparing *KO1/KO2* with *miR-1000 Rescue* and WT control flies using Kaplan-Meier statistics and log-Rank (Mantel-Cox) test.

### *miR-1000* regulates Neuropeptide-Like Precursor 1 (*Nplp1*) in the central nervous system

*Drosophila miR-1000* was first identified as a neuroprotective miRNA in the CNS (Verma et al. 2015). Despite being only 2% of the total body mass, the brain consumes ∼25% of total energy intake; hence, metabolism plays a crucial role in maintaining proper brain function. Accumulating evidence has suggested a strong association between obesity, metabolic dysfunction, and neurodegeneration (Procaccini et al. 2016; Uranga and Keller 2019; de et al. 2021). However, the underlying molecular mechanisms remain unknown. Prior analysis of *miR-1000* function in the CNS demonstrated its role in regulating the expression of *VGlut* (Vesicular Glutamate Transporter) and protecting against neurodegeneration caused by excitotoxic glutamatergic signaling (Verma et al. 2015). Thus, we investigated whether excess *VGlut* in *miR-1000* mutants contributes to the observed metabolic phenotypes. To test this, we reduced *VGlut* levels in *miR-1000* mutants by expressing *VGlut-RNAi* in *miR-1000* expressing neurons using a *miR-1000-GAL4* (*miR-1000-GAL4* generated by switching the miniwhite gene with *GAL4* in *KO2* mutants (Verma et al. 2015) and subjected the flies to starvation. No significant difference in total fat levels (S2_A) and their ability to survive starvation (S2_B) was observed between *miR-1000* mutants and *miR-1000* mutants with reduced *VGlut* levels, even though *VGlut* knockdown in these neurons was sufficient to rescue locomotor and neurodegeneration phenotypes in *miR-1000* mutants (Verma et al. 2015). Since a single miRNA can regulate multiple biological processes by targeting more than one target gene, we searched for additional target genes that might be relevant to the *miR-1000*-mediated metabolic phenotypes. To identify these targets, we quantified the transcript levels in the brain for the top 10 CNS-specific targets predicted by the TargetScanFly (https://www.targetscan.org/fly_72/) program. Besides *VGlut*, we observed a modest yet consistent elevation in the transcript level of a neuropeptide *Nplp1* (Neuropeptide-like precursor 1) encoded by gene *CG3441* in *miR-1000* mutants using RT-qPCR analysis of head tissue (Fig. 2A).

**Figure 2:**
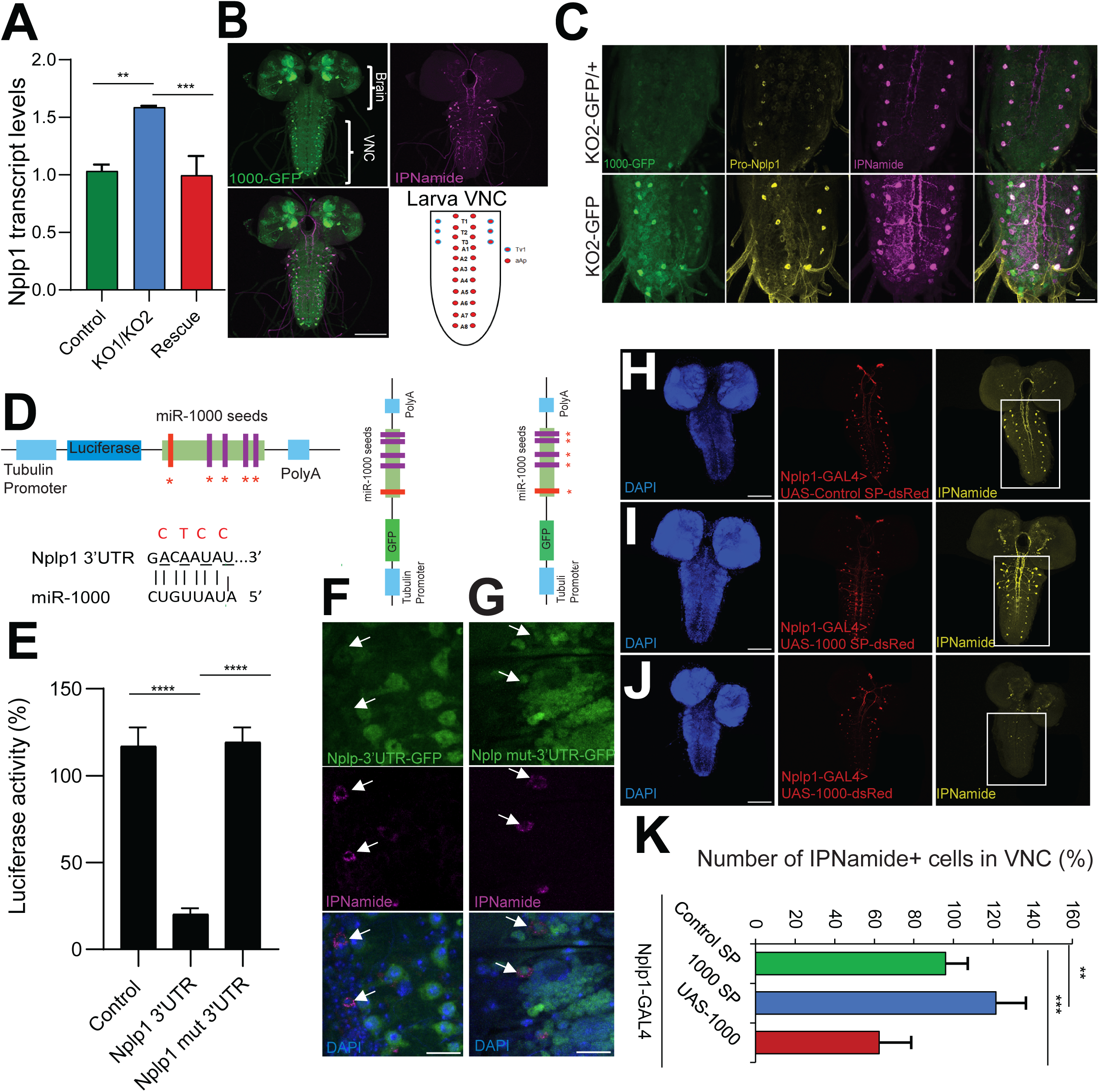
*miR-1000* regulates *Nplp1* in the Central nervous system (CNS). A) RT-qPCR showing upregulation of *Nplp1* transcript level (normalized to rp49) in *miR-1000* mutants (*KO1/KO2*) compared to WT control and *miR-1000 Rescue*. Bars represent an average of three biological replicates, n= 20 flies each, error bars ±SD. p=0.0014 *KO1/KO2* vs WT control, p=0.001 *KO1/KO2* vs *miR-1000 Rescue* using one-way ANOVA with Tukey’s multiple comparisons. B) Larval CNS showing expression of *miR-1000* (green) using *miR-1000-GFP* knock-in and *Nplp1* in *miR-1000-GFP* heterozygous brains using its derivative peptide IPNamide (purple) antibody in the brain and Ventral nerve cord (VNC). Scale bar: 100μm. Schematic showing a pattern of *Nplp1* expression in Tv1 and dAp neurons in larval VNC. C) Larval ventral nerve cord (VNC) showing expression of *miR-1000* (green), Pro-Nplp1 (yellow), and IPNamide peptide (purple) in heterozygous (*KO2-GFP/+,* upper panels) and homozygous (*KO2-GFP,* lower panels) *miR-1000-GFP* mutants. Scale bar: 25μm. Pro-*Nplp1* and IPNamide are elevated in *miR-1000* homozygous mutants (*KO2-GFP*, lower panels) compared to heterozygous mutants. D) Schematics showing the design of luciferase reporter with *Nplp1 3’UTR* having five seeds or MRE (microRNA recognition element) of *miR-1000* driven by tubulin promoter. *Nplp1* 3’UTR contains five target seeds (four strong seeds (purple) and one weak seed (red) of *miR-1000*. Normal seed pairing of *Nplp1 3’UTR* with *miR-1000*. Normal seed pairing is shown in black, and the mutated seed sequence in the *Nplp1 3’UTR* is shown in red. E) Luciferase reporter assay showing direct regulation of *Nplp1 3’UTR* by *miR-1000* in S2 cells. *Nplp1 3’UTR* showed a significant reduction (p<0.0001) in luciferase activity upon transfection with *miR-1000*, which was rescued by mutating all five seed sequences in *Nplp1 3’UTR* (p<0.0001) using two-tailed unpaired Student’s T-test. Bars represent an average of four biological replicates, error bars ±SD. F) Schematics showing *Nplp1 3’UTR-GFP* reporter having five seeds of *miR-1000* driven by the tubulin promoter. Magnified confocal images from VNC showing expression of *Nplp1 3’UTR-GFP* reporter (green) and IPNamide (purple) in wild-type background. dAP neurons expressing IPNamide and marked by the arrow show a lack of *Nplp1-3’UTR-GFP* reporter expression. Scale bar: 15μm. G) Schematics showing *Nplp1 3’UTR-GFP* reporter having five mutated seeds of *miR-1000* (shown by red star) driven by the tubulin promoter. Magnified confocal images from VNC showing expression of *Nplp1 3’UTR-GFP* reporter (green) and IPNamide (purple). dAP neurons expressing IPNamide and marked by the arrow show the presence of *Nplp1-3’UTR-GFP* reporter expression upon mutation of *miR-1000* seeds. Scale bar: 15μm. H) I) J) Larval VNC expressing *miR-1000 SP* (Sponge) showing increased IPNamide expression (H, yellow, middle panel) and reduction in IPNamide expression upon expressing *UAS-miR-1000* (I, yellow, lower panel) compared to the *control SP* (J, yellow, upper panel) using a *Nplp1-GAL4*. Red indicates *Nplp1-GAL4* expression with *UAS-dsRed* associated with *Control SP, 1000 SP,* and *UAS-1000*, and nuclei are stained with DAPI (Blue). Scale bar: 100μm. K) Quantification of Tv1 and aAP cells expressing IPNamide in VNC of *Nplp1-GAL4>Control SP, Nplp1-GAL4>1000 SP,* and *Nplp1-GAL4>UAS-1000*. Bars represent an average of at least six biological replicates. Error bars ±SD. p-value=0.0038 *Nplp1-GAL4>Control SP* vs *Nplp1-GAL4>1000 SP* and p-value=0.0008 *Nplp1-GAL4>Control SP* vs *Nplp1-GAL4> UAS-1000* using one way ANOVA with Tukey’s multiple comparisons.

The biological function of *Nplp1*-derived neuropeptides has been largely unknown so far, but their exquisitely selective distribution in the CNS indicates a possible role in interneurons within both the brain and Ventral Nerve Cord (VNC). *Nplp1* encodes a single precursor pro-protein (Pro-NPLP1) that can be processed into multiple peptides (IPNamide, NPLP-1-VQQ, MTYamide, NAP, and NIATMARLQSAPSTHRDP) (Baggerman et al. 2002; Verleyen et al. 2004). To decipher the biological function of *Nplp1*, we first examined its expression in the larval and adult CNS. *Nplp1* neurons have been previously characterized as a specific marker of neuropeptidergic neuronal cell fates in the larval CNS (Stratmann and Thor 2017). Subpopulations of 28 *Nplp1*-expressing neurons have been previously mapped to thoracic ventrolateral Tv1 and dorsal medial dAp neurons in the larval ventral nerve cord (VNC) (Stratmann and Thor 2017) (Fig. 2B). However, their biological function in the larval or adult CNS remained unknown.

To track NPLP1 peptide expression in the CNS, we used Pro-NPLP1 (precursor of Nplp1) and IPNamide (a derivative peptide of Nplp1) antibodies (Verleyen et al. 2004). Additionally, we generated an *Nplp1-GAL4* driver by cloning the *Nplp1* upstream regulatory genomic portion next to *GAL4* to track its gene expression (see Methods). Neuropeptides are often expressed in neuronal clusters with complex axonal projections, which can obscure the individual neuron morphology. Therefore, a combination of neuropeptide antibodies along with *Nplp1-GAL4*-driven expression was used to identify and characterize a more complete spatial pattern of *Nplp1*-positive neurons. IPNamide displayed little overlap with *1000-GFP* (*1000-GFP* generated by switching the miniwhite gene with *GFP* in the *KO2* mutant) in both larval (Fig. 2B, S3_A, S3_B) and adult brains and the ventral nerve cord (VNC) (S3_G, S3_H), consistent with a mutually exclusive regulatory relationship between a miRNA and its target gene. Pro-NPLP1 and IPNamide showed mostly overlapping expression patterns in the larval (S3_C, S3_D) and adult CNS (S3_I). Our *Nplp1-GAL4* driver recapitulated *Nplp1* expression as expected from prior analysis of the cis-regulatory modules (CRM) promoter fragment used (Stratmann and Thor 2017) and showed overlapping expression with IPNamide, along with some additional adjacent neurons (S3_E, S3_F). These expression data indicate that the *Nplp1* transcript has much more widespread expression compared to the final derivative NPLP1 peptides in the CNS. *Nplp1*-derived peptides might have also been secreted from the *Nplp1-GAL4* expressing cells, which is likely why they exhibit a faint or no expression in cells where *Nplp1-GAL4* expression is still detectable. Since *Nplp1* neurons are well characterized in larval VNC, we examined changes in the expression of Pro-NPLP1 and derivative IPNamide in *Nplp1*-positive VNC neurons. We observed a significant increase in both Pro-NPLP1 and IPNamide levels in *miR-1000* mutants (Fig. 2C, lower panel) compared to heterozygous control VNCs (Fig. 2C, upper panel).

### Neuropeptide, *Nplp1,* is a direct target of *miR-1000* in the Central Nervous System (CNS)

To determine if *Nplp1* mRNA can be directly targeted by *miR-1000*, we constructed a luciferase reporter containing the wild-type *Nplp1 3’UTR* with 5 predicted *miR-1000* complementary seed sequences (4 strong seeds shown in purple, 1 weak seed shown in red in Fig. 2D, S4_A). The *Nplp1 3’UTR* showed strong repression of luciferase activity in *S2* cells upon *miR-1000* expression (Fig. 2E, S4_B), suggesting a direct targeting of *Nplp1 3’UTR* by *miR-1000*. To assess the contribution of individual *miR-1000* seeds in the *Nplp1 3’UTR*, we first mutated the three strongest seeds individually and in combination with each other. However, a significant derepression by individual mutated seeds could not be achieved (S4_B). Therefore, we mutated all 5 seeds (Fig. 2D, S4_C) to abolish *miR-1000* binding to the *Nplp1 3’UTR* completely. This resulted in complete derepression of luciferase activity (Fig. 2E). To further investigate the regulation of *Nplp1 3’UTR* by *miR-1000* in the CNS, we generated an *Nplp1-3’UTR-GFP* reporter driven by the ubiquitous tubulin promoter *in vivo*. We did not observe *GFP* expression in IPNamide-expressing neurons in the VNC in wild-type conditions (Fig. 2F). However, *GFP* expression was significantly restored in the *Nplp1-3’UTR-GFP* reporter when all five *miR-1000* seed sequences were mutated (Fig. 2G). Both luciferase and *GFP* reporter data demonstrate that *miR-1000* strongly and directly targets *Nplp1* transcripts through all five complementary seeds in its *3’UTR*.

To test the regulation of *Nplp1* by *miR-1000* specifically in *Nplp1* neurons, we inhibited *miR-1000* function specifically in larval *Nplp1* neurons using an *Nplp-GAL4* with *miR-1000 SP* (Sponge); *miR-1000 SP* contains 10 copies of complementary *miR-1000* binding sequences to inhibit its function. Both *miR-1000 SP* and *Scramble Control SP* constructs were marked with *DsRed*, allowing us to track the expression of *Nplp1-GAL4* in *miR-1000 SP* or *Control SP* flies by the presence of the *DsRed* fluorescent protein. We quantified the number of *Nplp1* neurons in the VNC samples with detectable expression of IPNamide. Previous reports suggest that *Nplp1* neurons are well-defined and fixed in number (28 neurons) in larval VNC (Stratmann and Thor 2017). Inhibition of *miR-1000* function in *Nplp1* neurons resulted in increased staining by IPNamide-specific antibody (Fig. 2I, rightmost panel), as well as an increase in the number of *Nplp1* neurons with detectable IPNamide expression (Fig. 2K) compared to *Control SP* (Fig. 2H, 2K). Conversely, overexpression of *miR-1000* in *Nplp1* neurons diminished the expression of IPNamide in the VNC (Fig. 2J, 2K). These data, combined with our analysis of *miR-1000* targeting of the *Nplp1 3’UTR* reporter assays, strongly suggest that *Nplp1* and its derivative peptides are regulated by *miR-1000*.

### Elevated levels of *Nplp1* are responsible for the metabolic phenotypes of *miR-1000* mutants

To test whether elevation in *Nplp1* levels contributes to the metabolic defects of *miR-1000* mutants, we genetically reduced *Nplp1* levels using a hypomorphic insertion mutant, *Nplp1^EY11089,^* in the *miR-1000* mutant background. The combination of the *miR-1000* mutant with the *Nplp1^EY11089^* allele significantly reduced body weight (Fig. 3A), fat levels (Fig. 3B) and decreased fly survival upon starvation compared to *miR-1000* mutants alone (Fig. 3C). To establish that this effect is coming selectively from *Nplp1* neurons, we knocked down *Nplp1* by expressing *Nplp1-RNAi* specifically in *Nplp1* neurons using *Nplp1-GAL4* in the *miR-1000* mutant background. We observed a significant reduction in body weight (Fig. 3A) and a nearly complete rescue of both fat levels (Fig. 3D) as well as starvation resistance (Fig. 3E) in flies expressing *Nplp1-GAL4>Nplp1-RNAi* in the *miR-1000* mutant background.

**Figure 3:**
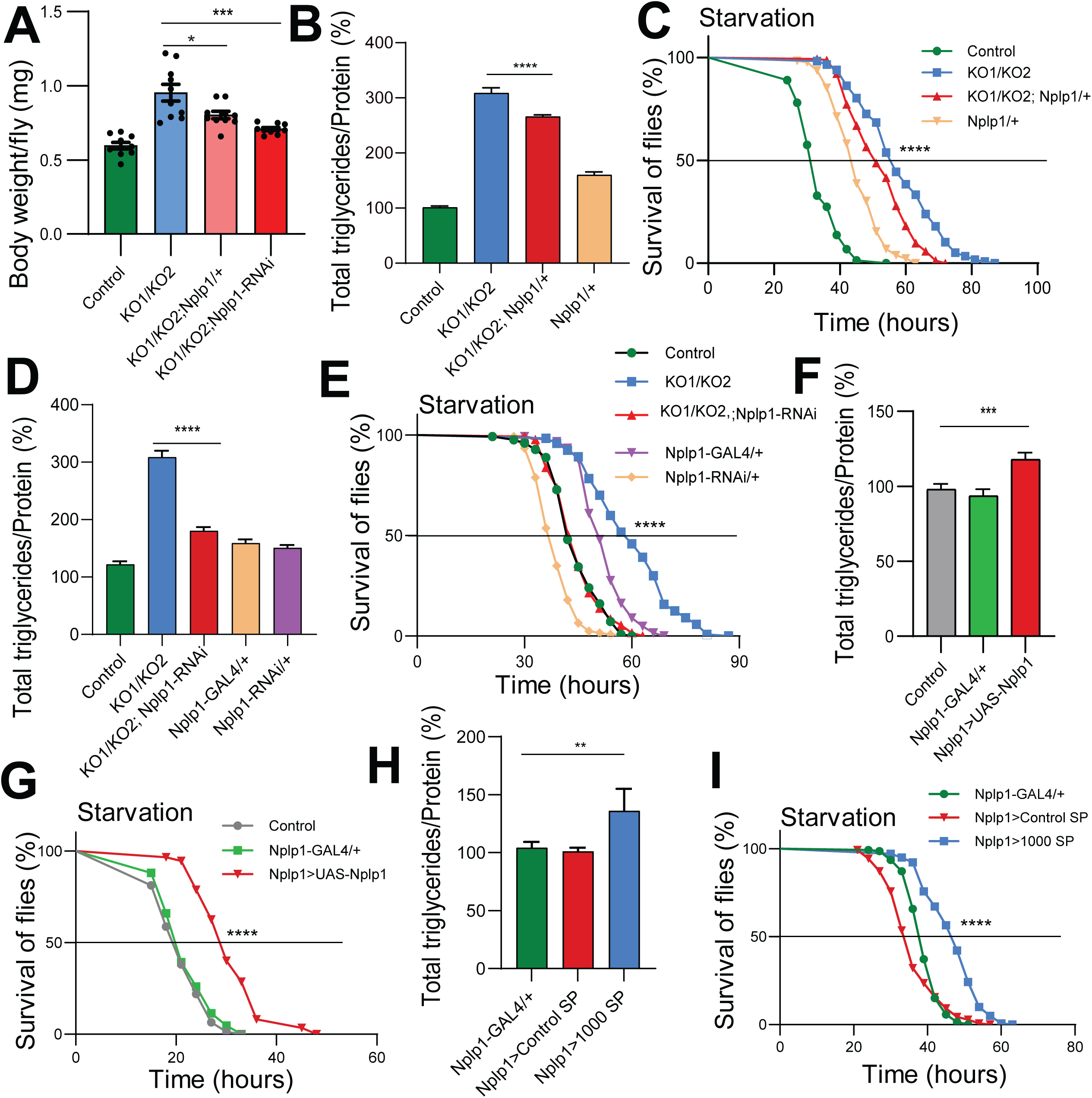
Elevated Nplp1 causes increased fat storage and starvation resistance. A) Body weight graph shows that reduction in *Nplp1* levels using *Nplp1^EY11089^*hypomorphic mutant (*KO1/KO2*; *Nplp1/+*, Pink, p= 0.0127) or expressing *Nplp1-GAL4>Nplp1-RNAi* in *miR-1000* mutant background (*KO1/KO2; Nplp1-RNAi*, red, p <0.0001) have reduced body weight compared to *miR-1000* mutant, *KO1/KO2* (blue) flies using one-way ANOVA with Tukey’s multiple comparisons. Bars represent an average of 5 biological replicates of 20 male flies each, error bars ±SEM. B) Percent total triglyceride levels (normalized to total protein). *Nplp1^EY11089^*hypomorphic mutant in *KO1/KO2* (*KO1/KO2; Nplp1/+*) reduces total triglycerides levels compared to *KO1/KO2* alone (p<0.0001). Wild type (WT), Control, and *Nplp1/+* are used as controls. Bars represent an average of four biological replicates, error bars ±SD. p<0.0001 comparing Controls with *KO1/KO2*, and *KO1/KO2* with *KO1/KO2; Nplp1* using one way ANOVA with Tukey’s multiple comparisons. C) Percent survival of flies upon starvation showing significant rescue of *KO1/KO2; Nplp1*/+ compared to *KO1/KO2*, p<0.0001 comparing *KO1/KO2* (n=117) with *KO1/KO2, Nplp1/+* (n=154), control (n=73) and *Nplp1/+* (n=129) using Kaplan-Meier statistics and log-Rank (Mantel-Cox) test. D) Percent total triglyceride levels showing reduction of triglyceride levels when *Nplp1-RNAi* is expressed in *Nplp1* neurons in *KO1/KO2* (p<0.0001). WT control, *Nplp1-GAL4/+,* and *Nplp1-RNAi/+* are used as controls. Bars represent an average of four biological replicates, error bars ±SD. p<0.0001 comparing control with *KO1/KO2, Nplp1>Nplp1-RNAi/+* with *KO1/KO2*, *KO1/KO2* compared to control, *Nplp1-GAL4/+* and *Nplp1-RNAi/+* using one way ANOVA with Tukey’s multiple comparisons. E) Percent survival of flies upon starvation showing significant rescue of *KO1/KO2; Nplp1>Nplp1-RNAi* compared to *KO1/KO2*, p<0.0001 comparing *KO1/KO2* (n=120) with *KO1/KO2, Nplp1>Nplp1-RNAi* (n=130), control (n=125), using Kaplan-Meier statistics and log-Rank (Mantel-Cox) test. F) Increased triglyceride levels in *Nplp1-GAL4>UAS-Nplp1 (Nplp1>UAS-Nplp1)* flies compared to *Nplp1-GAL4/+* and WT control. Bars represent an average of four biological replicates, error bars ±SD. p=0.001 control vs *Nplp1>UAS-Nplp1* and p=0.0002 *Nplp1-GAL4/+* vs *Nplp1>UAS-Nplp1* using one way ANOVA with Tukey’s multiple comparisons. G) Percent survival of flies upon starvation, showing significant starvation resistance in flies overexpressing *Nplp1* in *Nplp1* neurons. p<0.0001 comparing *Nplp1>UAS-Nplp1* (n=147) with control (n=155), and *Nplp1-GAL4/+* (n=150) using Kaplan-Meier statistics and log-Rank (Mantel-Cox) test. H) Increased triglyceride levels in flies where *miR-1000* is knocked down in *Nplp1* neurons. Bars represent an average of four biological replicates, error bars ±SD. p=0.0094 comparing *Nplp1>1000 SP* with *Nplp1-GAL4/+* and p=0.0054 comparing *Nplp1>1000 SP* with *Nplp1>Control SP* using one-way ANOVA with Tukey’s multiple comparisons. I) Increased survival of flies upon starvation where *miR-1000* is knocked down in *Nplp1* neurons. p<0.0001 comparing *Nplp1>1000 SP* (n=140) with *Nplp1-GAL4/+* (n=140) and *Nplp1>Control SP* (n=140) using Kaplan-Meier statistics and log-Rank (Mantel-Cox) test.

Conversely, to confirm that overexpression of *Nplp1* alone is sufficient to cause metabolic defects, we overexpressed *UAS-Nplp1* in *Nplp1* neurons using the *Nplp1-GAL4*. Indeed, we observed a significant increase in fat storage (Fig. 3F) and increased starvation resistance (Fig. 3G) in flies overexpressing *Nplp1*. This confirmed that overexpression of *Nplp1* is sufficient to cause metabolic defects. To prove that levels of *Nplp1* and subsequent metabolic phenotypes are controlled by *miR-1000* in *Nplp1* neurons, we depleted *miR-1000* using a *miR-1000 SP* specifically in *Nplp1* neurons. Knockdown of *miR-1000* specifically in *Nplp1* neurons caused flies to store more fat (Fig. 3H) and survive longer without food (Fig. 3I), hence confirming that *miR-1000*-mediated regulation of *Nplp1* specifically in *Nplp1* neurons is responsible for maintaining fat storage and survival upon food deprivation.

### Genetic manipulations of *Nplp1* also affect the overall lifespan of an organism

Metabolism plays a crucial role in survival and longevity (Owusu-Ansah and Perrimon 2014). Since *miR-1000* regulates both neurodegeneration and metabolism, it may represent a link between these two pathologies. In Humans, metabolic syndrome and neurodegenerative diseases both majorly affect middle-aged or elderly people. Indeed, obesity in middle age increases the risk of Alzheimer’s Disease by 2-fold (Whitmer et al. 2007). In *Drosophila*, *miR-1000* levels are also known to decrease with advancing age (Verma et al. 2015) and might increase the risk of metabolic and neurodegenerative dysfunction. Consistent with this idea, *miR-1000* mutants have been shown to have a short lifespan (S5_A). To investigate if *Nplp1* plays a role in determining the lifespan of *miR-1000* mutants, we examined the lifespan of flies where *Nplp1* levels were reduced in the *miR-1000* mutant background, either by using the *Nplp1^EY11089^* mutant allele (S5_B) or by expressing *Nplp1-RNAi* specifically in *Nplp1* neurons (S5_C). Both conditions significantly improved lifespan, suggesting that the metabolic defects caused by elevated *Nplp1* also contribute to the short lifespan of *miR-1000* mutants, and Nplp1-expressing neurons are also relevant to organismal survival.

### *miR-1000* regulates the size and density of lipid droplets (LDs) of fat bodies

Lipid droplets (LDs) are dynamic organelles that are responsible for lipid storage and mobilization according to the energy demands of an organism. In *Drosophila*, LDs within the fat body serve as the functional equivalent of mammalian liver and adipose tissue. They display distinct spatial organization, categorized as perinuclear (nLD) and peripheral LDs (pLD). Perinuclear lipid droplets (nLDs) are typically larger and are responsible for more stable lipid storage, are enriched in triacylglycerols (TAGs), and are less metabolically active. Peripheral lipid droplets (pLDs), found near the plasma membrane, are usually smaller and denser and are often associated with the process of lipolysis (lipid breakdown). These subpopulations differ in size, density, and likely function, reflecting local metabolic environments. Changes in LD morphology are often associated with imbalanced lipid dynamics and metabolic homeostasis (Olzmann and Carvalho 2019). To investigate this, we analyzed the size and density of pLDs and nLDs in *miR-1000* mutants using Nile Red lipid staining. In *miR-1000* mutants, pLDs were smaller (Fig. 4A, 4B) and denser (Fig. 4A, 4C) compared to WT controls and *miR-1000 Rescue,* indicating a highly dynamic and metabolically active state with active lipid turnover. Conversely, nLDs in *miR-1000* mutants were larger (Fig. 4D, 4E) and less dense (Fig. 4D, 4F) than WT controls and *miR-1000 Rescue*, suggesting a cellular state favoring long-term lipid storage over immediate energy mobilization. These findings were corroborated using a different lipid stain, BODIPY (S6_A, S6B). The co-existence of smaller, denser peripheral LDs and larger, less dense perinuclear LDs indicates a spatial separation of lipid function. This organization enables the organism to maintain energy homeostasis under fluctuating nutrient or stress conditions, reflecting a physiologically adaptive lipid metabolic mechanism.

**Figure 4:**
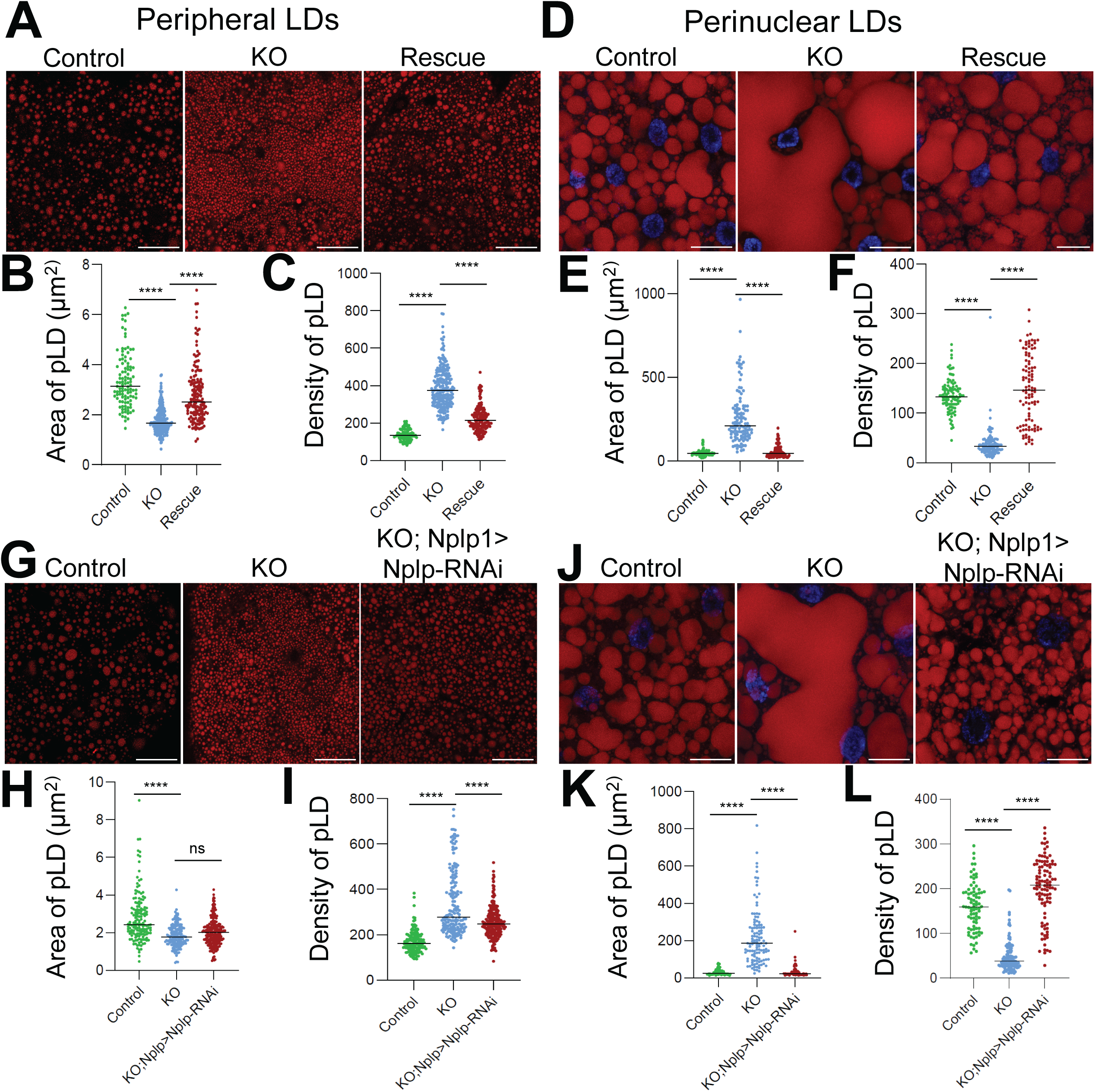
Loss of *miR-1000* affects lipid droplet (LD) size and density. A) Confocal images showing Peripheral lipid droplets (pLD) stained with Nile red present on the surface of larval fat body of WT control, *miR-1000 KO*, and *miR-1000 Rescue*. Scale bar: 25μm. B) Graph showing Area of pLD of larval fat bodies of WT control, *miR-1000 KO*, and *miR-1000 Rescue*. pLD of *miR-1000 KO* (n=226) is smaller as compared to WT Control (n=105, p<0.0001) and to *miR-1000 Rescue* (n=154, p<0.0001) using one-way ANOVA with Tukey’s multiple comparisons. C) Graph showing density of pLD of larval fat bodies of WT control*, miR-1000 KO*, and *miR-1000 Rescue*. pLD of *miR-1000 KO* (n=226) is denser as compared to WT Control (n=105, p<0.0001) and to *miR-1000 Rescue* (n=154, p<0.0001) using one-way ANOVA with Tukey’s multiple comparisons. D) Confocal images showing Nuclear lipid droplets (nLD) stained with Nile red present around the nucleus (marked by DAPI in Blue) of larval fat body of WT control, *miR-1000 KO*, and *miR-1000 Rescue*. Scale bar: 25μm. E) Graph showing Area of nLD of larval fat bodies of WT control, *miR-1000 KO*, and *miR-1000 Rescue*. pLD of *miR-1000 KO* (n=112) is bigger as compared to WT Control (n=103, p<0.0001) and to *miR-1000 Rescue* (n=99, p<0.0001) using one-way ANOVA with Tukey’s multiple comparisons. F) Graph showing density of nLD of larval fat bodies of WT control, *miR-1000 KO*, and *miR-1000 Rescue*. pLD of *miR-1000 KO* (n=112) is less dense as compared to WT Control (n=103, p<0.0001) and *miR-1000 Rescue* (n=99, p<0.0001) using one-way ANOVA with Tukey’s multiple comparisons. G) Confocal images showing pLD stained with Nile red present on the surface of larval fat body of WT control, *miR-1000 KO*, and with reduced levels of *Nplp1* using *Nplp-RNAi* expressing in *Nplp1* neurons in *miR-1000 KO* background (*KO; Nplp1-GAL4>Nplp1-RNAi*). Scale bar: 25μm. H) Graph showing Area of pLD of larval fat bodies of WT control, *miR-1000 KO*, and *miR-1000 KO; Nplp1-GAL4>Nplp1-RNAi*. pLD of *miR-1000 KO; Nplp1-GAL4>Nplp1-RNAi* (n=225) are smaller as compared to WT Control (n=154, p<0.0001) but are similar in area to *miR-1000 KO* (n=168, p=0.1036) using one-way ANOVA with Tukey’s multiple comparisons. I) Graph showing density of pLD of larval fat bodies of WT control, *miR-1000 KO*, and *miR-1000 KO; Nplp1-GAL4>Nplp1-RNAi*. *miR-1000 KO; Nplp1-GAL4>Nplp1-RNAi* (n=225) are less dense as compared to *miR-1000 KO* (n=168, p<0.0001) but denser compared to WT Control (n=154, p<0.0001), and using one-way ANOVA with Tukey’s multiple comparisons. J) Confocal images showing nLD stained with Nile red present around the nucleus (marked by DAPI in Blue) of larval fat body of WT control, *miR-1000 KO*, and *miR-1000 KO; Nplp1-GAL4>Nplp1-RNAi*. Scale bar: 25μm. K) Graph showing Area of nLD of larval fat bodies of WT control, *miR-1000 KO*, and *miR-1000 KO; Nplp1-GAL4>Nplp1-RNAi*. nLD of *miR-1000 KO* (n=115) are bigger as compared to *miR-1000 KO; Nplp1-GAL4>Nplp1-RNAi* (n=108, p<0.0001) and to WT Control (n=89, p<0.0001) using one-way ANOVA with Tukey’s multiple comparisons. L) Graph showing density of nLD of larval fat bodies of WT control, *miR-1000 KO*, and *miR-1000 KO; Nplp1-GAL4>Nplp1-RNAi.* nLD of *miR-1000 KO* (n=108) are less denser as compared to *miR-1000 KO; Nplp1-GAL4>Nplp1-RNAi*. (n=101, p<0.0001) but more dense than WT control (n=82, p<0.0001) using one-way ANOVA with Tukey’s multiple comparisons.

To further validate the contrasting energy states of pLDs and nLDs, we used a *Lsd-1-GFP* reporter expression and localization, which can provide critical insight into the metabolic state of the LDs. *Lsd-1 (Lipid storage droplet-1)* is the *Drosophila* homolog of mammalian *perilipin-1* (PLIN-1), an important lipid droplet surface protein that regulates the access of lipases to LDs (Beller et al. 2008). High *Lsd-1-GFP* expression was observed on both pLDs and nLDs in *miR-1000* mutants compared to WT controls (S7_A, S7_B), indicating a coordinated suppression of lipolysis by blocking access to the lipases and overall more stable lipid storage over mobilization. This suggests a cellular strategy to conserve lipids, favor storage, and maintain lipid droplet integrity, often seen during anabolic states or stress buffering in *Drosophila* (Kuhnlein 2012).

To test if *Nplp1* plays any role in this lipid storage dynamics, we checked LDs from fat bodies of the flies with reduced levels of *Nplp1* using *Nplp1-GAL>Nplp1-RNAi* in a *miR-1000* mutant background. Reducing *Nplp1* levels in *miR-1000* mutants resulted in no significant change in the size of pLDs (Fig. 4G, 4H), but a notable decrease in their density (Fig. 4G, 4I). This reduction in pLD density without a change in size suggests structural similarity but functional differences in the lipid droplets, potentially reflecting alterations in lipid composition or maturation. Furthermore, *miR-1000* mutants with reduced *Nplp1* levels exhibited a significant decrease in nLD size (Fig. 4J, 4K) and an increase in density (Fig. 4J, 4L), indicating a shift from lipid storage to increased lipolytic activity. Together, these findings reveal critical roles of *miR-1000* and *Nplp1* in maintaining lipid homeostasis and regulating the balance between lipid storage and mobilization.

### miR-1000 controls the triacylglycerol (TAG) synthesis and storage

To investigate alterations in lipid classes, fatty acid composition, and metabolic pathway activity regulated by *miR-1000* and *Nplp1*, we conducted lipidomics analysis (data available at https://www.iprox.cn/page/project.html?id=IPX0012162000) using *Drosophila* fat bodies from wild-type (WT) controls, *miR-1000* mutants, *miR-1000 Rescue*, and *miR-1000* mutants with reduced *Nplp1* levels (*KO; Nplp1-GAL4>Nplp-RNAi* is shown as *KO; Nplp1* in figure). Principal component analysis (PCA) of lipid profiles demonstrated distinct clustering among WT control, *miR-1000* KO, *miR-1000 Rescue* and *miR-1000* KO with reduced levels of *Nplp1 (KO; Nplp)* groups (Fig. 5A). The first two principal components accounted for 43.2% of total variance (PC1: 28.8%, PC2: 14.4%), indicating clear separation between genotypes. Specifically, WT controls and *miR-1000* mutants exhibited well-differentiated lipid profiles, whereas *miR-1000 Rescue* and *KO; Nplp1* groups displayed intermediate shifts away from *miR-1000* mutants, indicating substantial lipidomic remodeling. Comparative analysis revealed a significant elevation in triacylglycerides (TAGs) and reduction in phospholipids (e.g., phosphatidylcholines, phosphatidylethanolamines) in *miR-1000* mutant fat bodies compared to WT control (Fig. 5B). These findings suggest a metabolic shift favoring storage lipid accumulation, potentially at the expense of membrane synthesis or lipid droplet (LD) surface stability, which is critical for preventing LD fusion, which may cause nLDs to coalesce in *miR-1000* mutant. Pathway analysis further corroborated enhanced TAG synthesis and TAG storage (Fig. 5C), consistent with the observed increase in LD size in *miR-1000* mutants. Notably, both *miR-1000 Rescue* and *miR-1000* mutants with reduced levels of *Nplp1 (KO; Nplp)* showed a significant reduction in various TAG classes (Fig. 5D, 5E, S8_A, S8_B). These reductions in TAGs align with the restoration of LD size and density observed with Nile Red staining, compared to *miR-1000* mutants. Collectively, these results highlight the critical roles of *miR-1000* and *Nplp1* in regulating lipid metabolism and LD dynamics.

**Figure 5:**
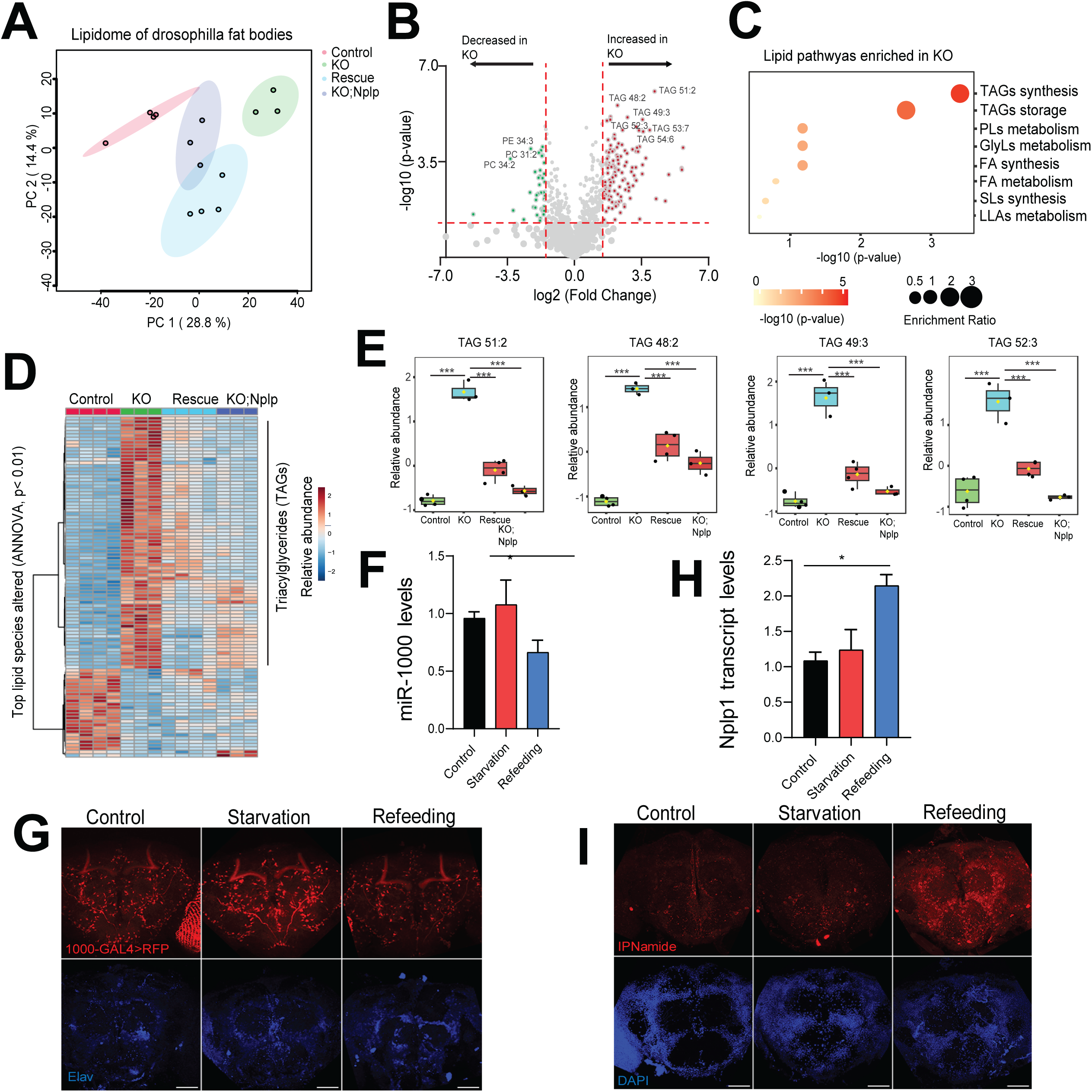
Lipidomics profiles of *miR-1000* mutants and changes in *miR-1000/Nplp1* expression with feeding. A) Principal component analysis (PCA) of the lipidome from WT control (pink), *miR-1000 KO* (green), *miR-1000 Rescue* (Blue), and *KO; Nplp1* (*KO; Nplp1-GAL4>Nplp1-RNAi*, purple) fat bodies. Data points indicate biological replicates. B) Volcano plot showing significantly altered serum lipid species in *miR-1000 KO* compared to WT control. Significance thresholds were set at p ≤ 0.05 (horizontal red line) and a fold change of ≥2 (vertical red line). C) Lipid pathways enriched in *miR-1000 KO* flies. D) Heatmap depicting top lipid species altered across the four groups, WT control, *miR-1000 KO, miR-1000 Rescue*, and *KO; Nplp1* (*KO; Nplp1-GAL4>Nplp1-RNAi*). The top lipid species altered were determined using one-way ANOVA, with a significant threshold at p ≤ 0.01. E) Relative abundance of TAGs in WT control, *miR-1000 KO*, *miR-1000 Rescue*, and *KO; Nplp1* (*KO; Nplp1-GAL4>Nplp1-RNAi*). p ≤ 0.001 when TAGs in *miR-1000 KO* are compared to WT Control, *miR-1000 Rescue* and *KO;Nplp1* (*KO;Nplp1-GAL4>Nplp1-RNAi*) one way ANOVA with Tukey’s multiple comparisons. F) RT-qPCR showing mature *miR-1000* levels in wild type control (0h starvation), Starvation (20h post-starvation), and Refeeding (3h after refeeding). p=0.57 comparing control with starvation, p=0.0358 comparing control with Refeeding. G) Adult brain images showing expression of *miR-1000* (red) using *miR-1000-GAL4* with *UAS-nRFP* in control (0h starvation), Starvation (20h post-starvation), and Refeeding (3h after refeeding). Elav (blue) is used to outline the brain neurons. Scale bar: 50μm. H) RT-qPCR showing *Nplp1* transcript levels in wild type control (0h starvation), Starvation (20h post-starvation), and Refeeding (3h after refeeding). p=0.585 comparing Control with Starvation, p=0.019 comparing Control with Refeeding. I) Adult brain of images showing expression of IPNamide (red) in Control (0h starvation), Starvation (20h post-starvation), and Refeeding (3h after refeeding). DAPI (blue) is used to outline the brain. Scale bar: 50μm.

### *miR-1000* and *Nplp1* levels change dynamically upon refeeding following starvation

Organisms frequently encounter periods of suboptimal food availability or starvation, particularly in the wild. These conditions trigger changes in signaling pathways and gene expression that are thought to facilitate adaptation to fluctuating food supplies (Fujikawa et al. 2009). Starvation also influences the expression and release of DILPs, as previously documented (Sudhakar et al. 2020). Consistent with these findings, we observed a significant reduction in *dilp2* transcript levels upon starvation (S9_A). To determine whether *miR-1000* levels respond to starvation, we assessed mature *miR-1000* levels at various time points (0h, 12h, 24h, 36h) during starvation in adult flies. Unlike insulin signaling, *miR-1000* levels remained unchanged during starvation (S9_B). However, a significant reduction in *miR-1000* levels was observed when flies were refed after starvation (Fig. 5F). These changes were further confirmed using a *miR-1000-GAL4* driver to express *UAS-RFP* in the adult brain (Fig. 5G). In parallel, refeeding after starvation led to a reciprocal increase in *Nplp1* transcript levels (Fig. 5H) and enhanced IPNamide expression (Fig. 5I). To investigate whether *miR-1000* levels respond to specific dietary components, we examined *miR-1000* expression following a high-sugar, a high-protein, and a high-fat diet. Interestingly, all dietary types triggered a reduction in *miR-1000* levels (S10_A). Together, our data suggest that maintenance of constant *miR-1000* levels might be essential for the dynamic mobilization of fat reserves during starvation, which confers a survival advantage to an organism during food deprivation. On the other hand, a reduction in *miR-1000* levels and a subsequent increase in *Nplp1* upon refeeding are required to send a signal to stop fat mobilization and switch to fat storage. These findings indicate that *miR-1000* and *Nplp1* play dynamic roles in re-establishing metabolic regulation following starvation.

Recent reports indicated that, similar to *miR-1000*, *miR-137* also regulates body weight and starvation resistance in *Drosophila* (Saedi et al. 2024). However, *miR-137* regulates metabolic homeostasis by targeting the *PTP61F* within the CNS. Both *miR-1000* and *miR-137* share an identical seed sequence for targeting mRNA gene transcripts (S11_A). To further investigate the relationship between *miR-1000* and *miR-137*, their expression patterns were examined in the CNS. The majority of *miR-1000* and *miR-137* expression did not overlap in larval CNS (S11_B) or adult brain (S11_C), although some overlap was observed in specific regions of the larval VNC (S11_B) and adult median brain regions (S11_C). A comparison of target genes identified by the TargetScan Fly program revealed only 126 common genes among the 717 predicted genes for *miR-1000* and 532 predicted genes for *miR-137* (S11_D). Notably, *miR-1000* and *miR-137* do not share *Nplp1* or *PTP61F* as common targets. However, *miR-137* has *Nplp3* as its target gene on the Targetscan list. *Nplp3* is a close family member of *Nplp1,* which is also expressed in the larval and adult CNS. These findings suggest that these miRNAs may regulate similar metabolic processes by targeting distinct members of the same gene family to maintain metabolic homeostasis.

Today, developed and developing countries face two contrasting problems. Developed nations face health problems like obesity and diabetes due to nutrition overabundance, whereas developing nations face malnutrition due to pest-infested crops and health problems due to insect-borne diseases like malaria. Recent studies reveal changes in mature *miR-1000* expression levels in the midgut of two *Anopheles* mosquito species after blood-feeding (Jain et al. 2014; Liu et al. 2017), reminiscent of our observations in *Drosophila*. Two *Nplp1*-family peptides show regulated expression dynamics immediately following blood-feeding of the insect *Rhodnius prolixus* (Sterkel et al. 2011), a vector of the trypanosome parasite that causes Chagas Disease in humans (Herbig-Sandreuter 1955). This reinforces our preliminary conclusion that the regulated expression of *Nplp1* is an important physiological mechanism in insect species. Since *Nplp1* is well conserved in all *Dipteran* insects, exploration of Nplp1-mediated feeding behavior might be useful for the development of biologically inspired insecticides.

In conclusion, the interplay between miRNAs and neuropeptides represents a sophisticated regulatory axis essential for maintaining metabolic homeostasis. Our findings reveal the critical role of *miR-1000* in modulating a specific neural circuit governed by *Nplp1*-expressing cells, which directly impacts body weight, fat storage, and survival under food deprivation. This study underscores the potential of miRNA-mediated neuropeptide regulation as a pivotal mechanism in metabolic control, paving the way for future research into its broader implications for health and disease.

## Acknowledgments

We thank the Bloomington *Drosophila* Stock Center (BDSC, Indiana) and the Vienna *Drosophila* Resource Center (VDRC, Vienna) for fly stocks. We are grateful to Dr. Stefan Thor for providing anti-Pro-NPLP1 and to Dr. Lilian Schoofs for the anti-IPNamide antibody. We are especially thankful to Prof. Stephen Cohen for his generous support in the initial phase of this work. We thank Kah Junn Tan for the injection of *UAS-1000 SP, Control SP, 1000-GFP, 1000-GAL4*, and *miR-1000 Rescue* constructs, and Best gene (www.thebestgene.com) for the injection of *Nplp1-GAL4* and *UAS-Nplp1* constructs. We thank Dr. Nilay Yapici for helpful discussions and Dr. Stephen Liberles, Dr. Marie Bao, Dr. Diana Ho, and Soohong Min for their insightful comments on the manuscript. We also thank Nikon Imaging Center (NIC) and Dr. Lisa Goodrich at HMS for access to the confocal imaging facility. PV and DVV were supported by NIH grants NS090994, NS119932, and NS123207.

## Author Contributions

PV conceived the research, performed all experiments and analyzed data, PD and NND analyzed lipidomics data, PV wrote the paper, and DVV edited the paper and acquired funding to complete the project.

## Materials and Methods

### Fly strains

*W^1118^* flies were used as a wild-type control unless otherwise indicated. All flies were grown on standard cornmeal fly food with a 12-12 hrs light-dark cycle with 60% humidity. *miR-1000* mutants (*KO1* and *KO2*), *miR-1000 Rescue, miR-1000 KO2-GAL4*, and *miR-1000-GFP* fly lines were generated as described in Verma et al, 2015. *UAS-miR-1000 SP* (Sponge), *UAS-miR-1000, Nplp-GAL4,* and *UAS-Nplp1* were generated as described below. *Nplp^EY11089^*(BL 20253), *UAS-CD8-RFP* (BL 32219), and *UAS-CD8-GFP* (BL 1537) flies were obtained from the Bloomington *Drosophila* Stock Center (BDSC), and *Nplp-RNAi* (VDRC 14035) flies from the Vienna *Drosophila* Resource Center (VDRC).

### Plasmids and transgenic flies generation

*UAS-miR-1000 SP* (sponge) transgenic flies were generated by cloning 10 copies of the mature miR-1000 sequence (ACTGCTGTGTGTAGACAATATCCGG) into the NotI-XbaI site in *pUAST-dsRed* vector and injected at 51D location on the second chromosome.

For cloning of *UAS-miR-1000* overexpression transgenic flies, 200bp of the genomic portion was amplified using the following primers: F:5’GGCCGCTGCGAAATAAATCACTCCAACTAA3’, R:5’TAGAGTTTTTCTCAGGTTTTCCTGTCC 3’ and cloned into the NotI-XbaI site of the *pUAST-DsRed* vector. These flies are available in the Bloomington stock center with stock numbers BL 60656 (on the 3^rd^ chromosome) and BL 60655 (on the 2^nd^ chromosome).

For generation of *Nplp1-GAL4*, 835bp upstream *Nplp1* regulatory genomic region was cloned upstream of GAL4 using primer F-5’ TGAAGCATATGTTTGATTCAAGGT 3’, R-5’ CGGCTCAACTGTTAAGTGAGTTTCGT 3’ in *pCasper-GAL4* vector.

*UAS-Nplp1* transgenic lines are made by amplifying the 1628 bp region using primers, F-5’ CGAACGAAACTCACTTAACAGTTG 3’, R-5’ ACATTACATTTGGGGGATTTACAC 3’, and cloning into the *pUAST-dsRed* vector.

### Quantitative RT-qPCR

Total RNA from 15-20 fly heads was extracted with miRNeasy tissue/cells advanced micro kit (Qiagen, catalog no: 217684). RNA from each sample was subjected to cDNA synthesis by TaqMan microRNA Reverse Transcription kit (Applied Biosystems, cat no: 4366597) and qPCR by Taqman Universal master mix II, with UNG (applied biosystems, cat no: 4440038) for mature *miR-1000* transcript (*dme-miR-1000*, cat no: 4440886, Assay ID: 243259_mat) and *miR-137* (*dme-miR-137*, Cat no: 4440886, Assay ID: 006442_mat). Data was normalized to *U14* (cat no: 4427975, Assay ID: 001750), *U27* (cat no: 4427975, Assay ID:001752), and *snoR422* (cat no: 4427975, Assay ID: 001742). qPCRs were carried out using a QuantStudio™ 7 Flex Real-Time PCR System (ThermoFisher Scientific) machine.

For *Nplp1* mRNA qRT-PCR, total RNA was extracted with miRNeasy tissue/cells advanced micro kit (Qiagen, catalog no: 217684). cDNA was synthesized by using SuperScript IV VILO master mix (Invitrogen, cat no: 11756050), and qPCR was done using Fast SYBR green master mix (Applied Biosystems, cat no: 4385616). Primers, *Nplp1* F-5’ AGCGATTCCTAGACACATCCAA 3’ and *Nplp1* R-5’ GACTCCATATACGCCTCGTCTG 3’, were used to detect *Nplp1* transcript and normalized to *Ribosomal Protein 49* (*rp49* F-5’GCTAAGCTGTCGCACAAA 3’ and R-5’TCCGGTGGGCAGCATGTG 3’).

### Immunocytochemistry and Imaging

Larva samples were fixed in 4% paraformaldehyde (PFA) for 20 minutes. Brains were washed 4 times with PBS+ 0.1% Triton-X (PBT) before blocking with 5% Higgs for 2 hrs at room temperature. Samples were kept in primary antibody overnight at 4^Ο^C. Adult brains were dissected in ice-cold phosphate-buffered saline (PBS) and fixed in 4% Paraformaldehyde (PFA) for 30-40 minutes. Brains were washed several times with PBS+ 0.1% Triton-X (PBT) before blocking with 5% Higgs overnight at 4^Ο^C and kept in primary antibody for 36-48 hours at 4^Ο^C. The following primary antibodies were used: chicken anti-Pro-NPLP1 (1:1000, gift from Dr. Stefan Thor), rabbit anti-IPNamide (1:500, gift from Dr. Liliane Schoofs), and chicken anti-GFP (1:1000, ab13970). Samples were washed 3-4 times with PBS+ 0.1% Triton-X (PBT). Alexa Flour secondary antibodies (Invitrogen) for anti-mouse, anti-rabbit, and anti-chicken were used in a 1:500 dilution along with DAPI (1:1000, ThermoFisher Scientific, cat no: 62248) for 2 hrs at room temperature or overnight at 4^Ο^C. Brains were washed several times in PBT before mounting in SlowFade Gold Antifade Reagent (Invitrogen, cat no: S36936).

Immunofluorescence images were collected using a Zeiss LSM 800 confocal microscope and processed using ImageJ software (https://imagej.net/) and Adobe Photoshop.

### Body weight measurement

For body weight measurement, 10 male or female flies of the desired genotypes at 7 days of age were collected and anesthetized. Each group was weighed on a fine weighing scale in a pre-calibrated Eppendorf tube. At least 10 groups of 10 flies were used for each experiment, and the average was taken to calculate the weight of each fly.

### Starvation and lifespan assay

For starvation and lifespan assays, desired crosses were set up in cages with apple juice plates. Apple juice plates were changed every 12 hours for the collection of embryos. 50 first instar larvae were collected and transferred to the food vials to avoid overcrowding and to give them standard growth conditions without competition. Adult flies were collected upon eclosion within 12 hours. Males and females were housed together and aged for at least 5 days to deplete their larval fat reserves before subjecting them to starvation assays. Flies were flipped into new food vials every other day.

For starvation assays, 15-20 male or female flies of 7-day age were separated and transferred to the 1% Agar for starvation experiments. The number of dead flies was counted every 3 hours. Every experiment was performed with at least three biological replicates, each containing 15-20 flies. Survival curves were plotted using PRISM (www.graphpad.com).

For lifespan assays, 20 male or female flies were collected upon eclosion and kept in separate vials. Flies were flipped onto new food every 3 days, and the number of dead flies was counted before transferring live flies to a new vial. Survival curves were plotted using PRISM (www.graphpad.com).

### Refeeding and differential diet feeding experiments

For refeeding, 7-day-old male flies were starved for 18-20 hours and fed on normal cornmeal starch food for 3 hours before collecting samples for RNA extraction or for dissecting adult brains for staining.

For differential diet food, 7-day-old male flies were starved for 18-20 hours and were fed for 3 hours with a high sugar diet containing 30% sucrose (Sigma, cat no# S0389) or a high protein diet containing 10% peptone (ThermoFisher Scientific, cat no# 211677) and 10% tryptone (ThermoFisher Scientific, cat no# 211705) or a high fat diet containing 30% coconut oil ((ThermoFisher Scientific, cat no# 365475000) in Formula 4-24 instant *Drosophila* medium (Carolina, Cat no# 173200). Flies fed for 3 hours on Formula 4-24 instant *Drosophila* medium after 18-20 hours of starvation were used as a control. All flies were raised under similar conditions of temperature and humidity. At least three biological replicates of 10 male flies from each experiment were collected for RNA extraction and qPCR.

### Triglyceride measurement

10-15 male flies were collected at 7 days of age and flash-frozen in liquid nitrogen. Male flies were chosen for the experiments to avoid having any effect on the metabolic activities of the flies due to egg production and hormonal fluctuations. The flies were homogenized in a homogenization buffer (0.05% Tween 20) with Pink RINO lysis kit (Next Advance, SKU: PINKR1) beads using a Bullet Blender 24 Gold bead homogenizer (Next Advance). Triglyceride levels for each sample, along with the standard curve, were measured using a Serum Triglyceride Determination Kit (Sigma, cat no: TR0100-1KT) as per the manufacturer’s instructions. Samples were incubated at 37 °C for 10 minutes before taking absorbance readings at 550nm. Total protein was determined using the Quick Start Bradford protein assay kit (Biorad, cat no: 5000201). Each sample was run for both the triglyceride and protein amounts simultaneously. Each triglyceride measurement was normalized to total protein levels. Each experiment was done with at least four biological replicates.

### Lipid measurement using Lipidomics

Fat bodies from 60-80 3^rd^ instar larvae were dissected in ice-cold PBS for each sample and were extracted in butanol/methanol (1:1) with 5 mM ammonium formate.

For lipidomics, chromatographic separation was conducted using Vanquish UHPLC (Thermo Scientific, Waltham, MA, USA) and an Acquity CSH C18 column (1.7 µm, 2.1 mm × 100 mm) with a guard column (1.7 µm, 2.1 mm × 5 mm) (Waters, Milford, MA). The mobile phase A was water: acetonitrile 40:60, and mobile phase B isopropanol: acetonitrile 90:10, both mobile phases containing 10 mM ammonium formate, and 0.1% formic acid. The elution gradient was 20% B from 0 to 3 min, 55% B at 7 min, 65% B at 15 min, 70% B at 21 min, 88% B at 23 min, 100% B at 24 min held until 26 min, and 20% B at 28 min and held until 30 min. The flow rate was 0.35 mL/min. The autosampler was at 4°C. The injection volume was 5 µL. Needle wash was performed between samples using dichloromethane: isopropanol: acetonitrile at 1: 1: 1. The mass spectrometry was an Exploris 480 orbitrap (Thermo Scientific, Waltham, MA, USA). The ion source was H-ESI. Spray voltage was 3500 V for positive ions, and 2500 V for negative ions. Sheath gas was set at 50 arbitrary units (Arb), auxiliary gas 15 Arb, sweep gas 1 Arb, ion transfer tube at 325 °C, and vaporizer at 350 °C. The scan mode was data-dependent (dd)-MS^2^, covering 150-1600 m/z in both positive and negative polarities. The precursor ion scan had a resolution of 60,000. For product ion scan, the resolution was 15,000, isolation width 1.0 m/z, and collision energy 25%.

Thermo Scientific LipidSearch software version 5.0 was used for lipid identification and quantitation. First, the product search mode was used to identify lipids based on the exact mass of the precursor ions and the MS^2^ mass spectra of the product ion scan. The precursor and product tolerance was 10 ppm. The absolute intensity threshold of precursor ions and the relative intensity threshold of product ions were set to 30000 and 1%, respectively. Next, the search results from all samples were aligned within a retention time tolerance of 0.25 min. The annotated lipids were then filtered to reduce false positives by only including the lipids with a total grade of A and B.

### Lipidomics Analysis

#### Data Filtering and Preprocessing

To ensure the reliability of the data analysis, lipid species that were non-informative, such as those with values close to the detection limit, were excluded (PMID: 23543913, PMID: 36572652, PMID: 38587201). A filtering process was implemented using a relative standard deviation (RSD) threshold of >25% and an interquartile range (IQR) threshold of 10% to remove such species. For parametric statistical evaluations, the data were treated as normally distributed with consistent variance and were adjusted to approximate a Gaussian probability distribution (PMID: 23543913, PMID: 36572652, PMID: 38587201). Additionally, auto-scaling (unit variance scaling) was performed by mean-centering the data and normalizing each lipid species by its standard deviation (PMID: 23543913).

#### Principal component analysis and Data Visualization

To analyze group patterns, principal component analysis (PCA) was conducted, and statistical significance was determined using PERMANOVA in MetaboAnalyst 6.0 (PMID: 38587201). Distributions were calculated based on Euclidean distance derived from principal components. Heatmaps were created in MetaboAnalyst 6.0 using Euclidean distance and Ward clustering to illustrate lipid profiles in different fly groups, focusing on the most differentially expressed lipid species. For comparisons across multiple fly groups, a one-way ANOVA was employed, with a false discovery rate (FDR) cutoff at 0.01 for identifying significant lipid species.

#### Lipid pathway analysis

Lipid pathway analysis was carried out using significantly altered lipid species in *KO* fly groups, using the MetaboAnalyst 6.0 lipid pathway analysis feature.

### 3’UTR luciferase Reporter assays

*Nplp1 3′UTR* Luciferase reporters were expressed under the control of the tubulin promoter in the *pCasper-UAS-Firefly* luciferase plasmid. *Nplp1* wild type *3’UTR* genomic region of size 1084bp was amplified using the following primers.

*Nplp1 wild type 3’UTR* F-5’-ATCGCCGTGTAATTCTAGAGCGGTCGTCTGTCCAATTAC-3’

*Nplp1 wild type 3’UTR* R-5’-GCTGCAGGTCGACCTCGAGAGAGCTTTACCGCAAGTACG-3’

For mutating the *miR-1000* seed sequences in the *Nplp1 3’UTR*, the following primers were used in combination with *Nplp1* wild type *3’UTR* F and *Nplp1* wild type *3’UTR* R primers.

Seed1 mutF-5’ GGCGTCTTTTATGTGTAAATGCCTACACCTGTTTCCACTAGGTA 3’

Seed1 mutR-5’ TACCTAGTGGAAACAGGTGTAGGCATTTACACATAAAAGACGCC 3’

Seed 2 mutF-5’ GTTAACAAACTGCGTCAAAGCCTACACAACTAAAGAAAATCTCGAT 3’

Seed 2 mutR-5’ ATCGAGATTTTCTTTAGTTGTGTAGGCTTTGACGCAGTTTGTTAAC 3’

Seed 3 mutF-5’ATACTTTAATGTATTCAAGCCTACACAAAAATATTGCCTACACAATATTG 3’

Seed3 mut R-5’CAATATTGTGTAGGCAATATTTTTGTGTAGGCTTGAATACATTAAAGTAT 3’

Seed 4 mut F-5’ TTACGAATACTAAATATCGCCTACACACTCTTTATGCAAATAACGC 3’

Seed 4 mut R-5’ GCGTTATTTGCATAAAGAGTGTGTAGGCGATATTTAGTATTCGTAA 3’

Seed 5 mut F-5’ CAAGCCTACACAAAAATATTGCCTACACAATATTGTGGGGTAT 3’

Seed 5 mut R-5’ ATACCCCACAATATTGTGTAGGCAATATTTTTGTGTAGGCTTG 3’

For the *miR-1000* expressing plasmid, *miR-1000* was expressed under the tubulin promoter from a plasmid containing a 200 bp genomic region, amplified with the primers, F-5’TGCGAAATAAATCACTCCAACTAA 3’ and R-5’GTTTTTCTCAGGTTTTCCTGTCC 3’.

*S2* cells were transfected in 24-well plates with 0.025 μg of the *firefly Nplp1*-luciferase reporter plasmid, 0.025 μg of a *Renilla* luciferase plasmid as a transfection control, and 0.25 μg of the *miR-1000* expression plasmid or empty vector. Transfections were performed in triplicate, and experiments were performed at least four times. Dual-luciferase assays (Promega) were performed 2.5 days after transfection according to the manufacturer’s protocol.

### Statistics

All experiments are performed at least three times, along with appropriate controls. The data represented are an average of multiple replicates, and the error bar represents ± SEM or ± SD. Student’s t-test (Two-tailed, unpaired, unequal variance) was used to compare the statistical significance between the two groups. For more than two groups, the comparison was done using one-way ANOVA with Tukey’s multiple comparisons. For survival experiments including starvation assays and longevity assays, data were analyzed using Kaplan-Meier statistics and the log-Rank (Mantel-Cox) test, which considers differences in survival at the beginning, at mid-point, and at the end to determine significance level. p-value>0.05 was considered statistically significant.

## Data Availability statement

All study data are included in the article and supplementary information. Flies used in this study are either available in the Fly stock centers or available upon request from the authors.

